# MMGB/SA Consensus Estimate of the Binding Free Energy Between the Novel Coronavirus Spike Protein to the Human ACE2 Receptor

**DOI:** 10.1101/2020.08.25.267625

**Authors:** Negin Forouzesh, Alexey V. Onufriev

## Abstract

The ability to estimate protein-protein binding free energy in a computationally efficient via a physics-based approach is beneficial to research focused on the mechanism of viruses binding to their target proteins. Implicit solvation methodology may be particularly useful in the early stages of such research, as it can offer valuable insights into the binding process, quickly. Here we evaluate the potential of the related molecular mechanics generalized Born surface area (MMGB/SA) approach to estimate the binding free energy Δ*G*_*bind*_ between the SARS-CoV-2 spike receptor-binding domain and the human ACE2 receptor. The calculations are based on a recent flavor of the generalized Born model, GBNSR6. Two estimates of Δ*G*_*bind*_ are performed: one based on standard bondi radii, and the other based on a newly developed set of atomic radii (OPT1), optimized specifically for protein-ligand binding. We take the average of the resulting two Δ*G*_*bind*_ values as the consensus estimate. For the well-studied Ras-Raf protein-protein complex, which has similar binding free energy to that of the SARS-CoV-2/ACE2 complex, the consensus Δ*G*_*bind*_ = −11.8 *±* 1 kcal/mol, vs. experimental −9.7 *±* 0.2 kcal/mol.

The consensus estimates for the SARS-CoV-2/ACE2 complex is Δ*G*_*bind*_ = −9.4 *±* 1.5 kcal/mol, which is in near quantitative agreement with experiment (−10.6 kcal/mol). The availability of a conceptually simple MMGB/SA-based protocol for analysis of the SARS-CoV-2 /ACE2 binding may be beneficial in light of the need to move forward fast.

## Introduction

Emerged as a global threat to human health, the SARS-CoV-2 virus that causes COVID-19 disease has been studied widely since the start of 2020. ^1^ Despite sequence and structure similarities with other viruses,^2^ no highly effective treatment option for the novel coronavirus is available. As of today, about 14 million people across the globe have tested positively for the virus, and around 600,000 have died of COVID-19. ^3^ This fast-growing pandemic highlights the role of computational structural biology and computer-aided drug design (CADD), which have the ability to accelerate the slow and expensive process of drug discovery. ^4^ In structure-based drug discovery, accuracy and speed of the binding free energy prediction of drug-like compounds (ligands) to target biomolecules plays a key role in virtual screening of drug candidates.^5–7^ Despite decades of research, efficient and accurate computational prediction of binding free energies is still a challenge.^8–12^

In theory, the binding free energy of a molecular system can be estimated directly from thermodynamic first principles.^13^ However, for any realistic molecular systems, *e*.*g*., the complex of interest in this work that is made of more than 12,000 atoms,^14^ approximations must be made to make the estimate computationally feasible. For example, alchemical methods,^15,16^ simulate changes in the free energy along a pathway that sometimes reflects non-physical properties or literally “alchemy”. The required sample points along the pathway are generated via Monte Carlo or Molecular Dynamics (MD) simulations. Some of the popular methods in this class are thermodynamic integration (TI) and free energy perturbations (FEP).^17^ However, these simulations are still computationally expensive, specifically when it comes to high throughput virtual screening of thousands of potential drugs.

Remarkably more efficient, end-point free energy methods ignore details of the complex to unbound state pathway and estimate free energy on an ensemble of snapshots representing the initial (complex) and the final (unbound) states only. These snapshots can be generated by an MD simulation. Molecular mechanics Poisson-Boltzmann surface area (MMPB/SA) and molecular mechanics generalized Born surface area (MMGB/SA)^18–20^ are arguably among the most popular methods in this category. They are often used in docking projects where a quick estimate of binding affinities is required. ^21^ Leading docking software, for instance, AutoDock Vina^22^ and DOCK,^23^ rank the feasible poses of a ligand in a binding pocket based on a scoring function in which binding affinity plays an important role. End-point free energies can improve the accuracy of these scoring functions on-the-fly. Recently, MMGB/SA was employed to improve the accuracy of AutoDock Vina and Dock in the Drug Design Data Resource (D3R) Grand Challenge 4 (GC4).^24^

While calculations based on practical implicit solvation models such as generalized Born (GB) are arguably not as accurate as corresponding estimates based on the best available explicit solvent models, the use of implicit solvent has an undeniable appeal. And not only of computational efficiency: reasoning about physical origins of observed effects is often much more transparent in an implicit than explicit solvent.^25–27^ That last advantage may be particularly valuable now, when so much about COVID-19 structure-infectivity relationship remains unknown.

In this work, we employ MMGB/SA implemented in AmberTools18^28^ for binding free energy calculation of the SARS-CoV-2 spike receptor-binding domain (SARS-CoV-2 S RBD) and the human ACE2 receptor (PDB ID:6m0j), see Fig. 1. Through the MMGB/SA approach, the absolute binding free energy of a complex is calculated as the sum of gas-phase energy, solvation free energy, and entropic contributions averaged over several snapshots extracted from the main MD trajectory. A grid-based surface GB model is used for estimating the polar component of solvation free energy, coupled with water and atomic radii introduced earlier.^29^ Human H-Ras and the Ras-binding domain of C-Raf1, so-called Ras-Raf complex, ^6,30^ is chosen as the reference for the initial evaluation of the MMGB/SA model. Final results are compared with those from a few available relevant studies. The main goal of the work is a quick assessment of the potential of the simple and efficient MMGB/SA method to future studies of SARS-CoV-2 to ACE2 binding.

**Figure 1:**
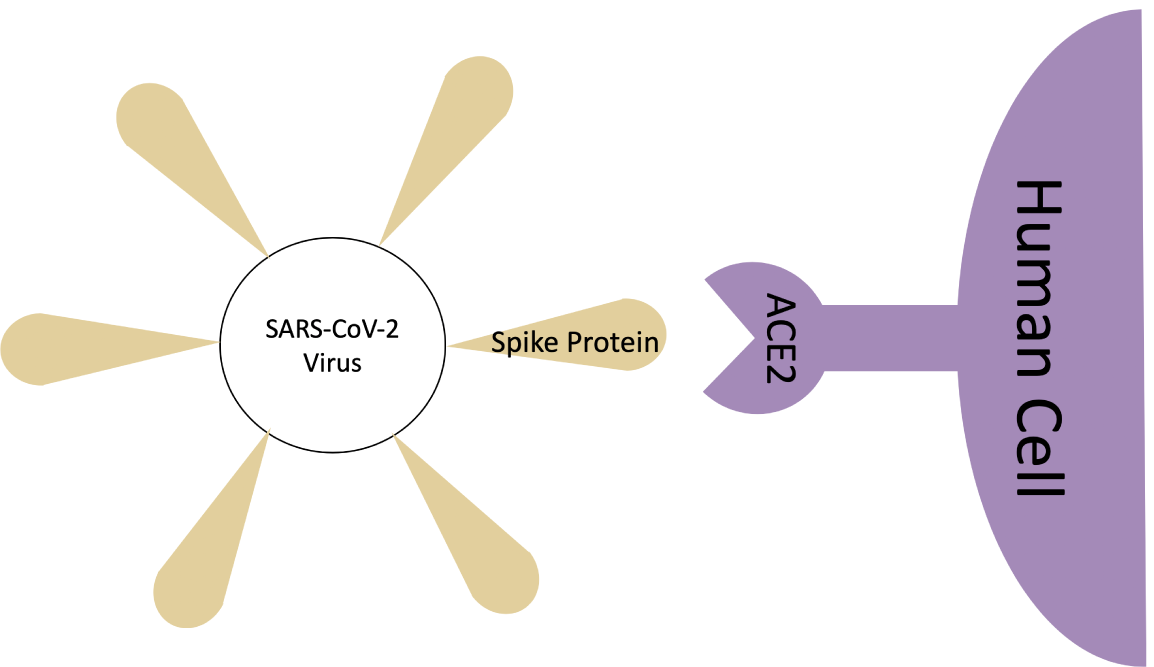
Binding scheme of the SARS-CoV-2 spike protein to the ACE2 human receptor.

## Methods and Materials

### Binding Free Energy Decomposition

Binding free energy, Δ*G*_*bind*_, of a molecular system is calculated as follows:

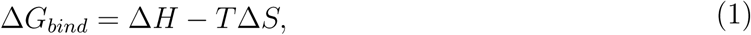

where Δ*H* is the enthalpy change of the system, *T* is the absolute temperature in K, and Δ*S* is the entropy change of the system. A high-level illustration of Δ*G*_*bind*_ between bound and unbound states of a solvated complex is shown in Fig. 2.

**Figure 2:**
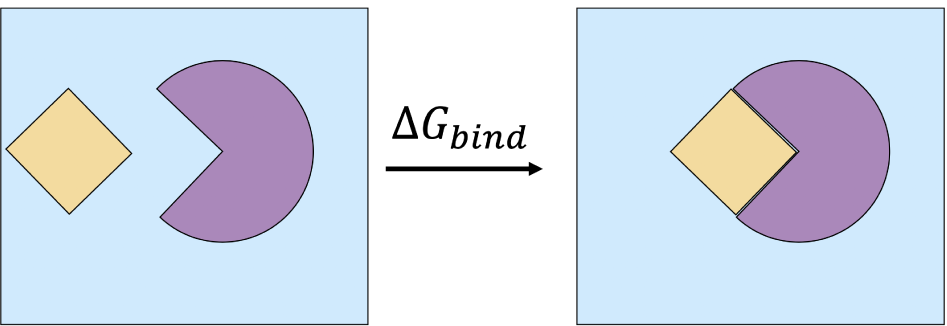
Binding a ligand (shown in yellow) to a protein receptor (shown in purple) in a box of solvent (shown in blue) releases binding free energy of Δ*G*_*bind*_. A negative sign o Δ*G*_*bind*_ indicates that spontaneous binding occurs, the magnitude of |Δ*G*_*bind*_| characterizes the binding strength (affinity).

In theoretical/computational studies, a useful way of calculating Δ*G*_*bind*_ is through a thermodynamic cycle shown in Fig. 3.

**Figure 3:**
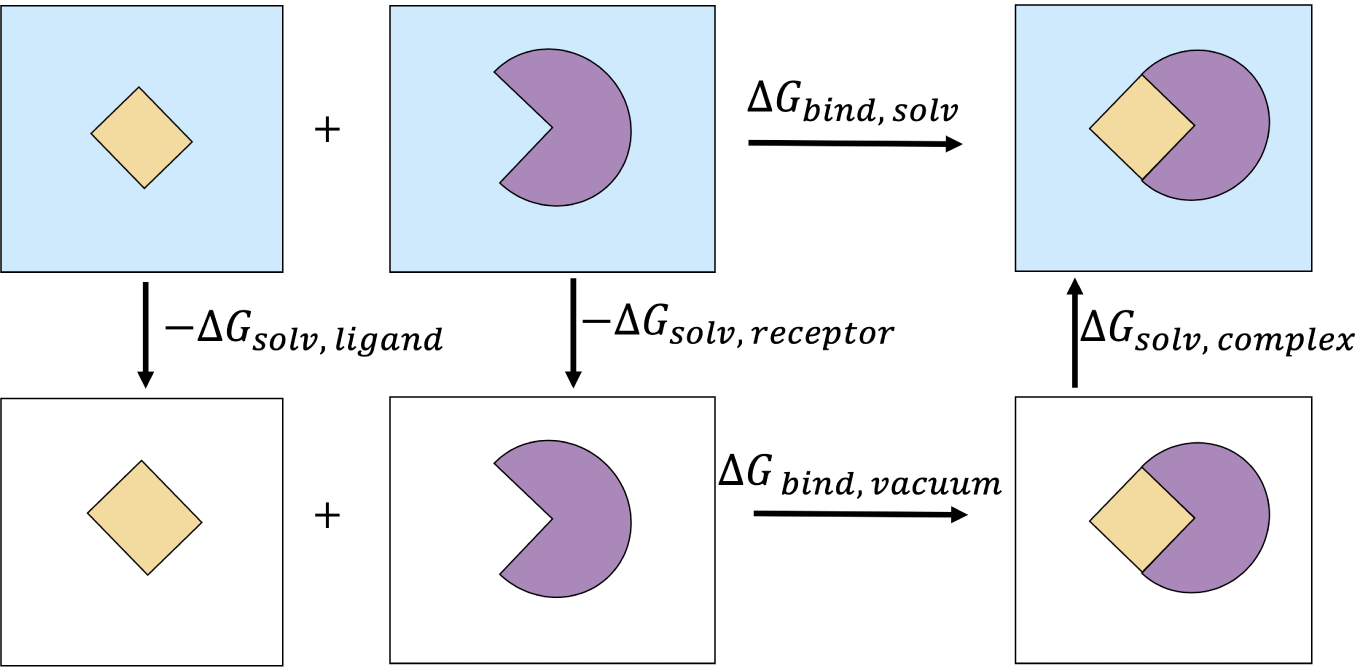
The thermodynamic cycle used here to estimate the binding free energy of a protein-ligand complex in the solvent.

With this approach, Δ*G*_*bind,solv*_ is calculated as follows:

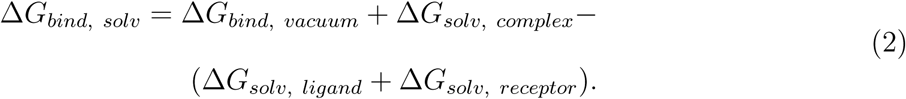

The solvation free energy, Δ*G*_*solv*_, is broken into the polar and non-polar components:

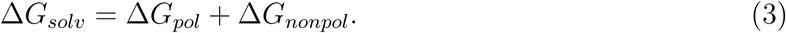

The free energy in vacuum, Δ*G*_*vacuum*_, is decomposed into the gas-phase energy (Δ*E*_*MM*_) and the configurational entropy of the solute (*T* Δ*S*):

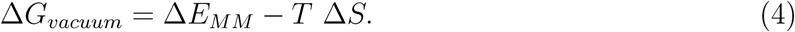

Note that the *T* Δ*S* above does not exactly correspond to *T* Δ*S* in Eq. 1; specifically, the entropy of solvent re-arrangement^25,27^ is subsumed into Δ*G*_*solv*_, see below, which is then considered a part of Δ*H*. Combining the free energy components defined above, we obtain: Δ*H* = Δ*E*_*MM*_ + Δ*G*_*pol*_ + Δ*G*_*nonpol*_. Our approaches for calculating Δ*G*_*solv*_, Δ*E*_*MM*_ and *T* Δ*S* are explained below.

### MMPB/SA Free Energy Methodology

MMPB/SA is a popular end-point free energy method which estimates Δ*G*_*solv*_ by Poisson-Boltzmann implicit solvent model, ^31^ while components of Δ*E*_*MM*_ are estimated based on a classical Molecular Mechanics force-field. (In MMGB/SA, discussed below and used here, the role of the PB is played by the faster GB). Significantly faster than the conventional Alchemical methods, MMPB/SA can be very useful, particularly in the early stages of structure-based virtual screening. As another important advantage, is that through MMPB/SA it is possible to decompose the total free energy into sub-components and measure their contributions separately. ^6,30^ This feature is certainly useful when it comes to comparing several different free energy methods. Finally, MMPB/SA is applicable to a wide range of structures, ^32^ from small host-guest systems to large protein-protein complexes with thousands of atoms.^6^

Through the MMPB/SA approach, the average of Δ*G*_*solv*_ is calculated on a collection of snapshots extracted from an MD simulation. Several decisions have to be made in applying the approach in practice. First, the computational protocol must be selected between the “single-trajectory” (one trajectory of the complex), or “separate-trajectory” (three separate trajectories of the complex, receptor and ligand). In this study, we choose the former protocol as it was shown^33^ to be not only much faster than the alternative, but also less “noisy” due to the cancellation of inter-molecular energy contributions. This protocol applies to cases where significant structural changes upon binding are not expected. Shown in Fig. 4, the single-trajectory MMPB/SA starts with the initial structure of the complex in vacuum. After solvating the structure in a solvent model, an MD simulation is performed to generate the snapshots for further analysis. Then, a relatively large number (typically *N >* 100) of uncorrelated snapshots are extracted to represent the structural ensemble. Next, binding free energies of these structures are calculated in the implicit solvent after removing the explicit solvent molecules. The average binding free energy over these snapshots is reported as the final Δ*G*_*bind*_.

**Figure 4:**
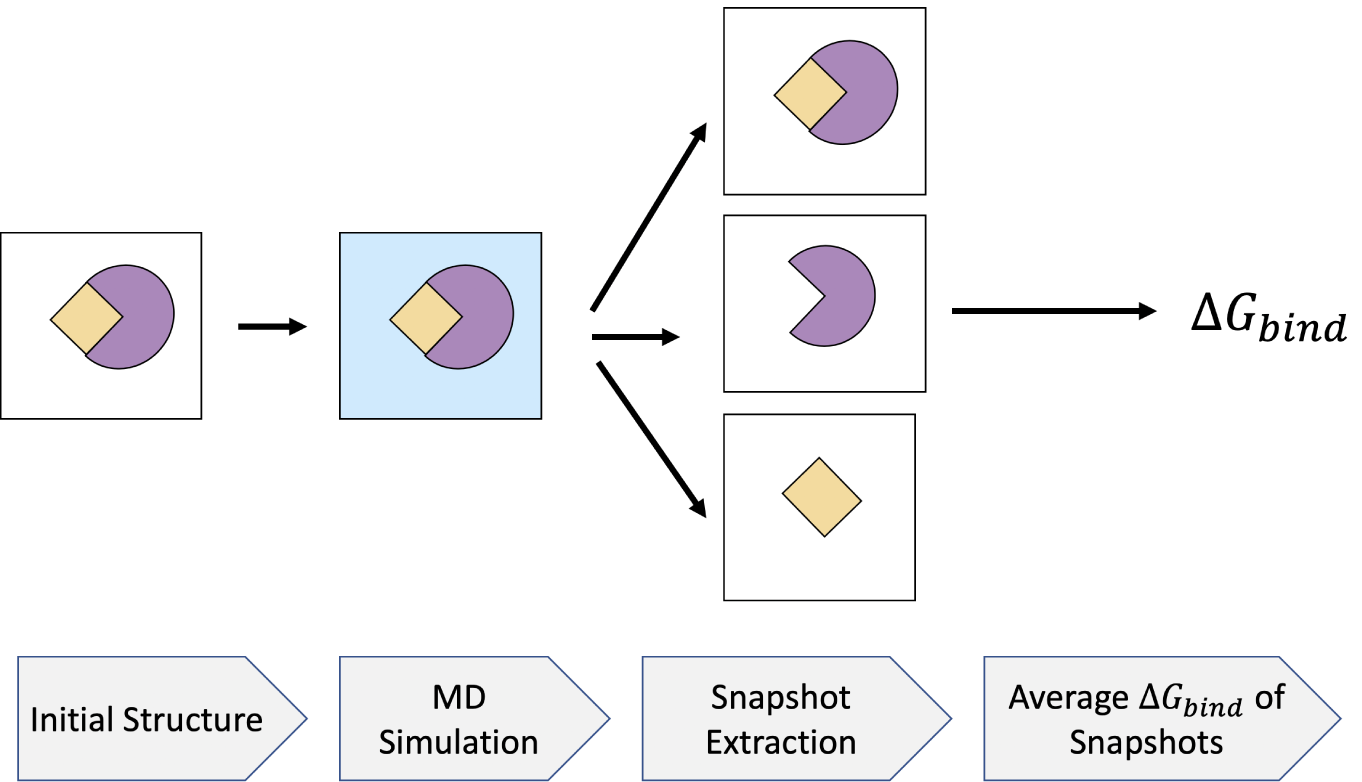
MMPB/SA flowchart. The initial structure of the complex is solvated using a water model. An MD simulation is run from which a relatively large number of snapshots are extracted. The average binding free energy of the snapshots is assigned as the binding free energy of the system.

With the single-trajectory protocol, the binding free energy of a protein-protein complex is formally calculated as follows:

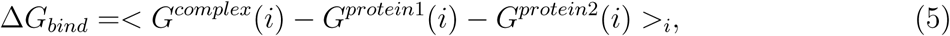

where *<* … *>*_*i*_ denotes an average over *i* snapshots extracted from the main MD trajectory. The implementation of this protocol is available in AmberTools18 in Perl^33^ and Python. ^34^

In this work the former is used to maintain consistency with the reference study^30^ opted for tuning the MMGB/SA model.

### Solvation Free Energy

#### Polar Component

A computationally efficient alternative to the PB, the GB implicit solvent model^35,36^ can be used for computing Δ*G*_*solv*_. Generally speaking, GB models have shown to be computationally less expensive than the PB models, although the deterioration of the accuracy has always been a concern. Here, we employed a grid-based surface GB model called GBNSR6, ^37^ which, in a recent study, ^38^ was shown to be the most accurate among several GB models in terms of the ability to approximate Δ*G*_*pol*_ relative to the numerical PB. In this work, Δ*G*_*pol*_ is calculated with the ALPB modification^39,40^ of the generalized Born^41^ model:

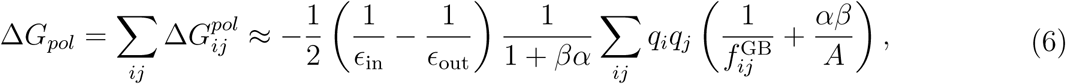

where *ϵ*_in_ = 1 and *ϵ*_out_ = 80 are the dielectric constants of the solute and the solvent, respectively, *β* = *ϵ*_in_*/ϵ*_out_, *α* = 0.571412, and *A* is the electrostatic size of the molecule, which is essentially the overall size of the structure that can be computed analytically. Here we employ the most widely used functional form 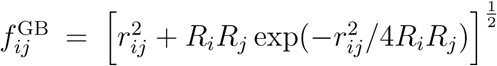, where *r*_*ij*_ is the distance between atomic charges *q*_*i*_ and *q*_*j*_, and *R*_*i*_, *R*_*j*_ are the so-called *effective Born radii* of atoms *i* and *j*, which represent each atom’s degree of burial within the solute. The effective Born radii, *R*, are calculated by the “*R*^6^” equation: ^42,43^

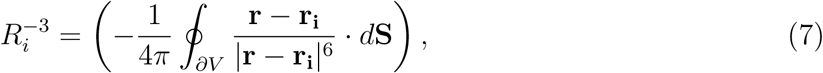

where ∂*V* represents the chosen representation of the dielectric boundary of the molecule, *d***S** is the infinitesimal surface element vector, **r**_**i**_ is the position of atom *i*, and **r** represents the position of the infinitesimal surface element. Uniform offset to the inverse effective radii is set to the default (optimal) value that is 0.028 Å^−1^.^44^ The screening effect of monovalent salt is introduced into Eq. 6 as is standard for the GB model;^35^ in our MMGB/SA calculations the salt concentration was set to 0.1 M.

#### Non-polar Component

A common method to estimate the non-polar contribution to the solvation free energy in Eq. 3 is to assume that it is proportional to the solvent accessible surface area (SASA) of the molecule:

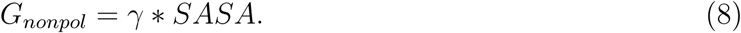

While there are more accurate methods to estimate the non-polar^45^ contribution, here we use the simple Eq. 8 for the sake of simplicity and consistency with ref. ^30^ Also for consistency with the same work, here we use *γ* = 0.0072 *kcal/mol/A*^2^. Atomic radii that form SASA not only play an important role in the non-polar component, but also enter the polar component through the dielectric boundary. Therefore, the right choice of atomic radii is crucial to the accuracy of binding free energy estimation. Three sets of atomic radii are used here: OPT1,^29,46^ bondi, and mbondi2. The first two are listed in Tab. 1. Mbonid2 is indeed bondi whose hydrogen atoms bound to a nitrogen are expanded from 1.2 Å to 1.3 Å, see .^47^ Carbon (C), hydrogen (H), oxygen (O), nitrogen (N) and sulfur (S) are the main atomic types in this study. The water probe radius is fixed to 1.4 Å.

#### Gas-Phase Energy

Gas-phase energy of the solute, Δ*E*_*MM*_, is the summation of internal energies, electrostatic energies, and van der Waals energies. In all of the MMGB/SA calculations reported here, Δ*E*_*MM*_ is calculated using the ff99 AMBER force field. The choice of this old force field is deliberate, and was initially motivated to ensure maximum consistency with, ^6^ which provides a very detailed analysis of MMGB/SA performance on Ras-Raf. Good agreement with experiment, Table 2, motivated us to use the same ff99 force field for all the subsequent MMGB/SA calculations reported here. All of the enthalpy calculations in this study are averages over 500 snapshots extracted from the main MD trajectory.

**Table 1:** Two sets of atomic radii in Å used in this study.

| | $\rho_C$ | $\rho_H$ | $\rho_N$ | $\rho_O$ | $\rho_S$ |
| --- | --- | --- | --- | --- | --- |
| bondi | 1.70 | 1.20 | 1.55 | 1.52 | 1.80 |
| OPT1 | 1.40 | 1.55 | 2.35 | 1.28 | 1.80 |

**Table 2:** MMGB/SA results on RAS-RAF. Means and the standard errors of the mean are listed. All the components are in *kcal/mol*. Consensus Δ*G*_*bind*_ = (Δ*G*_*bind*_(*bondi*) + Δ*G*_*bind*_(*OPT* 1))*/*2 For comparison, an earlier MMGB/SA estimate is also listed (it does not contribute to the consensus estimate). The experimental value is from isothermal titration calorimetry. ^52^ An offset of 1.79 *kcal/mol* has been subtracted from the Δ*H* component of MGB-based estimate for consistency with the author’s recommendation in. ^30^

|  | GBNSR6<br>(bondi) | GBNSR6<br>(OPT1) | **Consensus** | MGB<br>(bondi)<br>ref<sup>30</sup> | Exp. |
| --- | --- | --- | --- | --- | --- |
| $E_{MM}$ | $-937.42 \pm 2.07$ | $-937.42 \pm 2.07$ | | $-1308.6 \pm 2.5$ | |
| $\Delta G_{nonpol}$ | $-9.63 \pm 0.02$ | $-11.02 \pm 0.02$ | | $-9.5 \pm 0$ | |
| $\Delta G_{pol}$ | $887.39 \pm 2.06$ | $897.59 \pm 2.15$ | | $1275.3 \pm 2.4$ | |
| $\Delta H$ | $-59.66 \pm 0.27$ | $-50.85 \pm 0.43$ | | $-42.7 \pm 0.3$ | |
| $-T\Delta S$ | $43.39 \pm 0.98$ | $43.39 \pm 0.98$ | | $41.4 \pm 1.6$ | |
| $\Delta G_{bind}$ | $-16.27 \pm 1.01$ | $-7.46 \pm 1.07$ | **-11.8 ± 1** | $-3.1 \pm 1.6$ | $-9.7 \pm 0.2$ |

### Configurational Entropy

Normal-mode analysis (NMA) and quasi-harmonic analysis are of the two common methods for calculating configurational entropy of the solute. ^33^ Since the latter has shown poor convergence in several cases, NMA is selected for entropy calculations. ^30^ The main drawback of this method is the computational cost that becomes intractable for large systems, e.g., systems with more than 8,000 atoms in MMGB/SA (Perl version) of AMBER18 are not supported for NMA. To tackle this problem, one approach is to truncate the complex so that the binding interface is preserved in its original shape.^48^ In this study, the SARS-CoV-2 S RBD and ACE2 complex is truncated for NMA feasible calculations. An offset of 1.92 *kcal/mol* has been subtracted from the −*T* Δ*S* component of GBNSR6 calculations to address the concentration-dependency of the translational entropy at 1*M*, see ^30^ for details. NMA entropy calculations are done over 150 snapshots extracted from the main MD trajectory.

### Structure Preparation

#### Ras-Raf Complex

This well-studied complex was selected as the reference for testing the parameters of the MM/GBSA model. We used `tleap` module in AMBER18 to set up the input coordinate and topology files. The structure was solvated in a box of TIP3P^49^ water model (10 Å buffer). This choice of old water model and ff99 AMBER force field was deliberate, to ensure full consistency with. ^6^ The GTP molecule and the magnesium ion (*Mg*^2+^) were eliminated for the sake of simplification. Since the net charge was 0, no counterion was added.

#### SARS-CoV-2 S RBD and ACE2 Complex: Full Structure

H++ server^50^ was employed to protonate the complex at pH=7.5. The server automatically generates the solvated structure in a box of OPC^51^ explicit water model (10 Å buffer), with AMBER ff14SB force field. This full structure is used only for enthalpy calculations that are compared with those of the truncated complex structure for justifying the truncation approach, see below.

#### SARS-CoV-2 S RBD and ACE2 Complex: Truncated Structure

To execute NMA entropy calculation the original structure (PDB ID:6m0j) was truncated from 12,515 atoms (791 residues) to 7,286 atoms (463 residues) by removing residues, one by one, starting from the N-terminus of the spike protein, and the C-terminus of the ACE2 protein. The goal was to have fewer than 8,000 atoms remaining, while preserving sequence continuity of the resulting structure to facilitate the set up of MD simulations, see Fig. 5. The remaining atoms are still within 8 Å from the binding interface. The same protocol used for the full structure was employed for parameterization and solvation.

**Figure 5:**
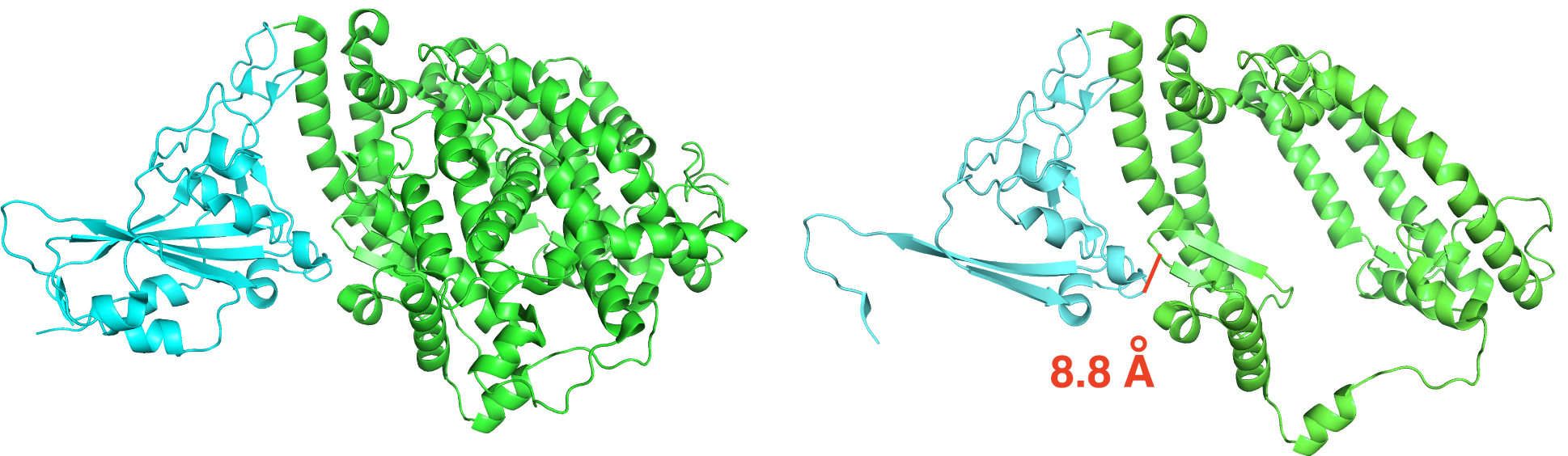
Truncation of SARS-CoV-2 S RBD used in the entropy estimate. The spike protein is in cyan, and the ACE2 receptor is in green. Left: original complex. Right: truncated complex. A pair of atoms on the binding interface that are 8.8 Å apart is shown in a solid red segment to illustrate the length scale.

### Trajectory Generation

All of the MMGB/SA estimates are based on snapshots extracted from MD trajectories generated as described below. The structures were prepared and equilibrated as follows. The solvated complexes were first energy minimized (max. minimization cycle of 1000), followed by 50 ps of heating (from 1 K to 300 K) at constant volume, followed by 50 ps of density equilibration at 300 K at constant 1 bar pressure, followed by another 2 ns of constant (1 bar) pressure equilibration at 300K. In these stages, atomic coordinates were restrained to their initial positions with 2 *kcal/mol/A*^2^. All simulations, including the production runs described below, were executed with the GPU-enabled pmemd.cuda MD engine in AMBER18, using Langevin dynamics with a collision frequency of 2*ps*^−1^ and an integration time step of 2 fs while the bonds involving hydrogen atoms were constrained by the SHAKE algorithm. Electrostatic interactions were approximated via the Particle Mesh Ewald (PME) method, with a non-bond cutoff set to 9 Å. Coordinates were recorded every 10 ps.

#### Ras-Raf Complex

A production of 10 ns was performed using the Ras-Raf structure prepared with the protocol described in Sec..

#### SARS-CoV-2 S RBD and ACE2 Complex: Full Structure

A production of 50 ns was carried out using the full structure described in Sec..

#### SARS-CoV-2 S RBD and ACE2 Complex: Truncated Structure

The same protocol used for the full structure was employed. A weak restraint of 0.01 *kcal/mol/A*^2^ was applied to the atoms of the truncated complex, relative to the X-ray positions, during the 50 ns production to prevent the truncated complex from falling apart. This restraint diminishes the discrepancy between the force filed and water model in the structure used for MD simulation (OPC, ff14SB) and the one for Δ*G*_*bind*_ calculations (TIP3P, ff99).

## Results and Discussion

### MM/GBSA on Ras-Raf

Here, we study the accuracy of Δ*G*_*bind*_ calculation using a different GB model (GBNSR6) coupled with two sets of atomic radii. According to Tab. 2, it is observed that Δ*G*_*bind*_ calculated by GBNSR6 with OPT1 radii underestimates the binding affinity whereas GBNSR6 coupled with bondi radii overestimates it. Yet, both of these results have better agreement with the experiment^52^ compared to the reference MGB model in .^30^ We have also noticed that a *consensus* estimate Δ*G*_*bind*_ = (Δ*G*_*bind*_(*bondi*) + Δ*G*_*bind*_(*OPT* 1))*/*2 is only within 2 kcal/mol off the experimental reference. Encouraged by this reasonable agreement with experiment, we have decided to utilize GBNSR6 with both bondi and OPT1 radii to produce a consensus estimate of Δ*G*_*bind*_ for the SARS-CoV-2 S RBD and ACE2 complex.

### MM/GBSA on the Truncated SARS-CoV-2 S RBD and ACE2

The RMSD of the truncated SARS-CoV-2 S RBD and ACE2 backbone compared to the crystal structure of the full complex is shown in Fig. 6. The trajectory is stable after 50 ns of production, with the RMSD from the X-ray reference of around 3.15 Å. Comparing the estimated Δ*H* between the truncated and original structures (results not shown) demonstrates that the relatively small difference between the two (about 1 kcal/mol using mbondi2 radii and 8 kcal/mol using OPT1 radii) indirectly validates the use of the truncation procedure. Shown in Tab. 3, our final estimates are presented, and compared to a previous computational study^53^ and experiment. These findings are briefly discussed in Conclusion.

**Figure 6:**
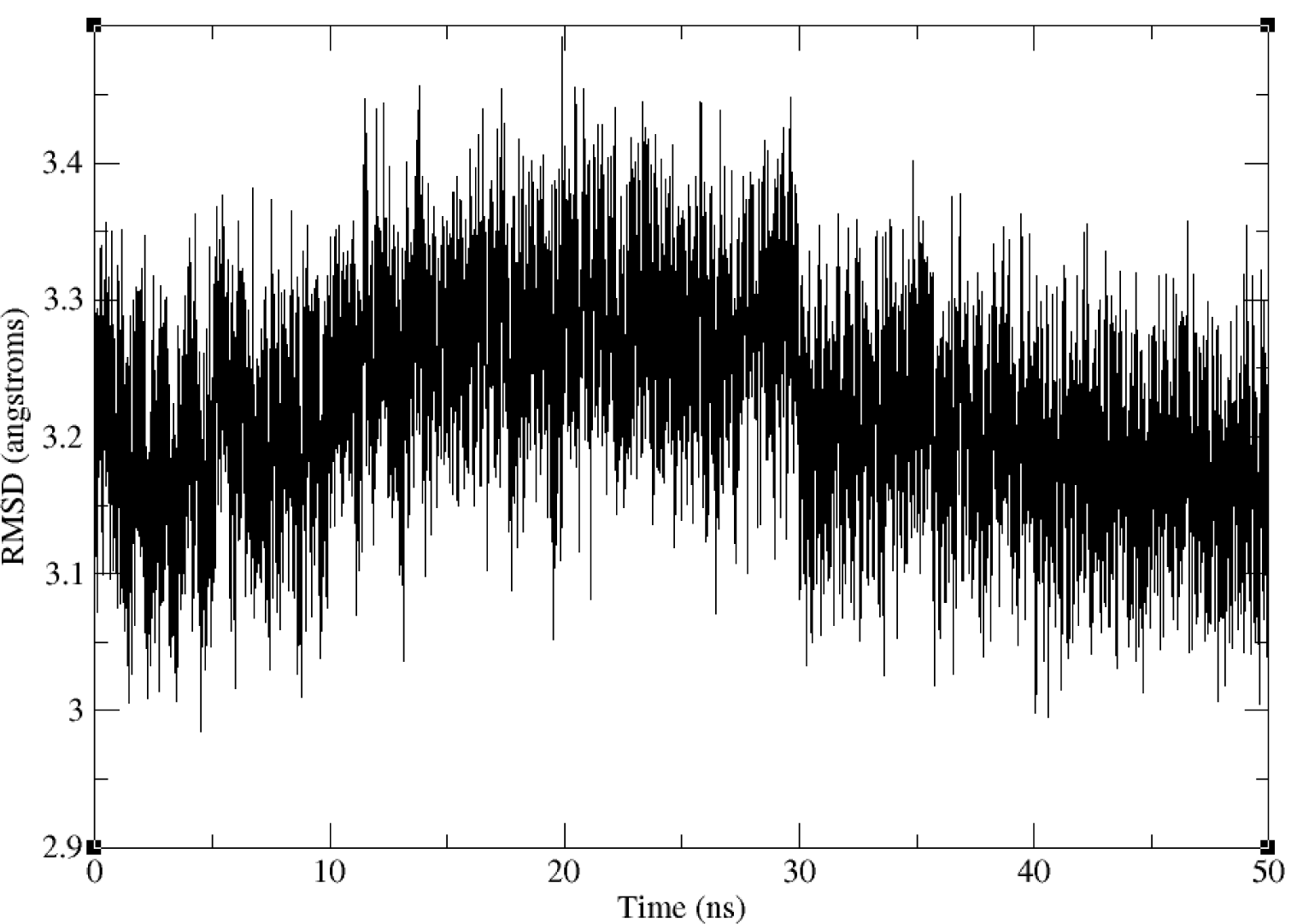
Backbone RMSD of the truncated SARS-CoV-2 S RBD and ACE2 complex, relative to the truncated part of the experimental crystal structure of the full complex, along the 50 ns production trajectory.

**Table 3:** MMGB/SA results on the truncated SARS-CoV-2 S RBD and ACE2 complex. Means and the standard errors of the mean are listed. All the components are in *kcal/mol*. Consensus Δ*G*_*bind*_ = (Δ*G*_*bind*_(*bondi*) + Δ*G*_*bind*_(*OPT* 1))*/*2 For comparison, a recently published MMGB/SA estimate is also listed (it does not contribute to the consensus estimate). Experimental value derived from a fit to surface plasmon resonance sensogram. ^54^

| | GBNSR6<br>(mbondi2) | GBNSR6<br>(OPT1) | Consensus | $GB^{OBC}$<br>(mbondi2)<br>ref<sup>53</sup> | Exp. |
| --- | --- | --- | --- | --- | --- |
| $E_{MM}$ | $-453.76 \pm 0.87$ | $-453.76 \pm 0.87$ | | $-761.06 \pm 3.96$ | |
| $\Delta G_{nonpol}$ | $-14.71 \pm 0.02$ | $-16.35 \pm 0.02$ | | $-12.21 \pm 0.06$ | |
| $\Delta G_{pol}$ | $401.55 \pm 0.81$ | $413.72 \pm 0.87$ | | $737.98 \pm 3.86$ | |
| $\Delta H$ | $-66.93 \pm 0.29$ | $-56.39 \pm 0.37$ | | $-35.30 \pm 0.60$ | |
| $-T\Delta S$ | $52.28 \pm 1.49$ | $52.28 \pm 1.49$ | | $13.56 \pm 0.70$ | |
| $\Delta G_{bind}$ | $-14.65 \pm 1.52$ | $-4.11 \pm 1.54$ | **-9.4 $\pm 1.5$** | $-21.74 \pm 0.65$ | **-10.6** |

### Entropy Convergence of the Truncated SARS-CoV-2 S RBD and ACE2 Complex

Entropy calculation is one of the most challenging and time-consuming parts of Δ*G*_*bind*_ estimation. In order to maintain the consistent protocol, 150 snapshots were selected for −*T* Δ*S* calculations. In addition, we conducted an investigation to examine whether a fewer number of snapshots would suffice to lead to a similar −*T* Δ*S*. A subset of 15 and 50 equidistant snapshots were collected from the set of 150 snapshots. According to Tab. 4 it is observed that −*T* Δ*S* calculated on 15 and 50 sample snapshots leads to a similar −*T* Δ*S* calculated on the whole set. Naturally, the standard error of the mean decreases as the sample size increases. However, this increase doesn’t affect the stability of the mean around 52 *kcal/mol*. Given entropy calculation as the bottleneck of Δ*G*_*bind*_ estimation, this observation suggests that with a relatively small set of snapshots it is possible to compute −*T* Δ*S* quite accurately.

**Table 4:**
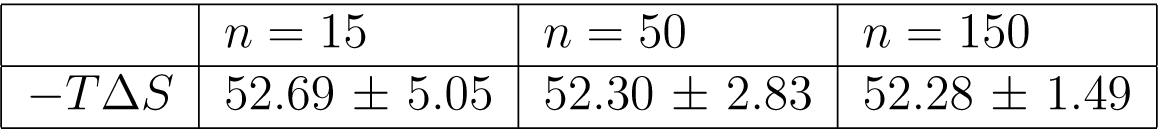
Entropy convergence of the truncated SARS-CoV-2 S RBD and ACE2 complex. Means and standard error of the means are listed. Increasing the number of equidistant snapshots *n* from 15 to 150 shows the stability of the entropy around 52 *kcal/mol*.

## Conclusion

In this study, we have evaluated the performance of a relatively new GB model, GBNSR6, and the newly introduced intrinsic atomic radii, in calculations of binding free energy of a well-studied protein-protein complex, Ras-Raf and of SARS-CoV-2 S RBD and ACE2 complex, critical in the mechanism of the novel coronavirus infection. Unlike many previous efforts, the new radii were specifically optimized to best reproduce the explicit solvent results particularly in the implicit solvent-based binding MMGB/SA estimates. We also employed the common bondi radii for the same calculations.

A better agreement with experiment of the absolute binding free energy for Ras-Raf, compared to previous work, was achieved for both radii sets; however each individual estimate either under- or over-estimated the experimental binding energy. We have therefore proposed a consensus estimate, which is the average of the two: for Ras-Raf the consensus based on OPT1 and bondi is in near quantitative agreement with experiment.

We applied the same approach to estimate Δ*G*_*bind*_ of the SARS-CoV-2 S RBD and ACE2 complex. As in the Ras-Raf case, the under- and over-estimation by each radii sets nearly cancelled, resulting in a consensus estimate essentially within 1 kcal/mol from the experimental reference. That essentially quantitative agreement paves the way for further exploration of SARS-CoV-2 S RBD and ACE2 complex with MMGB/SA, which has a number of advantages. Specifically, the MMGB/SA approach could be reasonably accurate for future analysis of *relative* binding free energies in this system, including the effects of mutations, relative contributions from various residues to Δ*G*_*bind*_, congeneric series of ligands, etc. Equally importantly, MMGB/SA is well-suited for reasoning for physical reasoning. Needless to say, additional thorough investigation is needed to see if the proposed approach can be extended to other complexes.

Both of the two separate estimates of Δ*G*_*bind*_ based on either bondi or OPT1 radii result in the expected near cancellation of the relatively large Δ*H* and −*T* Δ*S* terms, suggesting that each of these estimates “makes sense” on its own. Moreover, note that for both of the protein-protein complexes studied here, the two radii sets provide a fairly narrow range within which the experimental value lies. Specifically, Δ*G*_*bind*_ calculated with bondi radii is underestimated (too negative), whereas Δ*G*_*bind*_ calculated by OPT1 radii is over-estimated. One rationale for this behavior, which is the basis of the proposed consensus approach, could be that bondi and OPT1 radii sets have very different physical foundations behind them (geometry for the former and global optimization of the electrostatics for the latter), so the resulting errors in the corresponding electrostatic estimates are not as strongly correlated as for radii derived on the same principle.

All of the MD trajectories generated in this work are available from the authors upon request.

## Acknowledgment

The authors thank Dr. Holger Gohlke and Dr. Nadine Homeyer for their assistance with developing GBNSR6 in AMBER and their consultation. This work was supported by the NIH R21 GM131228.

## Notes

### Competing Interest Statement

The authors have declared no competing interest.

